# Pyrazinamide action is driven by the cell envelope stress response in *Mycobacterium tuberculosis*

**DOI:** 10.1101/2021.02.17.431758

**Authors:** Joshua M. Thiede, Nicholas A. Dillon, Michael D. Howe, Ranee Aflakpui, Samuel J. Modlin, Sven E. Hoffner, Faramarz Valafar, Yusuke Minato, Anthony D. Baughn

**Author notes:** Joshua M. Thiede and Nicholas A. Dillon contributed equally to this work. Correspondence and requests for materials should be addressed to Anthony D. Baughn.

## Abstract

Pyrazinamide (PZA) plays a crucial role in first-line tuberculosis drug therapy. Unlike other antimicrobial agents, PZA is only active against *Mycobacterium tuberculosis* at low pH. The basis for this conditional drug susceptibility remains undefined. In this study, we utilized a genome-wide approach to interrogate potentiation of PZA action. We find that mutations in numerous genes involved in central metabolism as well as cell envelope maintenance and stress response are associated with PZA resistance. Further, we demonstrate that constitutive activation of the cell envelope stress response can drive PZA susceptibility independent of environmental pH. Consequently, treatment with peptidoglycan synthesis inhibitors, such as beta-lactams and D-cycloserine, potentiate PZA action through triggering this response. These findings illuminate a regulatory mechanism for conditional PZA susceptibility and reveals new avenues for enhancing potency of this important drug through targeting activation of the cell envelope stress response.

## INTRODUCTION

*Mycobacterium tuberculosis*, the etiological agent of tuberculosis (TB), is the leading cause of death by a single pathogen, killing 1.5 million people each year^1^. Current TB short-course therapy consists of a four-drug regimen including isoniazid (INH), rifampicin (RIF), ethambutol (EMB) and pyrazinamide (PZA). Due to the unique activity of PZA against persistent populations of bacilli, its inclusion in therapy has led to a reduction in treatment times from 9 to 6 months and has dramatically reduced disease relapse rates^2^. Unfortunately, the increasing prevalence of drug resistant TB infections compromises the viability of effective treatment regimens. In 2015, the estimated global incidence of PZA resistance was 16%, accounting for as many as 1.4 million cases^3^. At least 70% of clinical PZA resistance can be attributed to loss-of-function mutations in *pncA* which encodes an amidase that is essential for conversion of the drug to its active form pyrazinoic acid (POA)^4^. With the expansion of deep sequencing based approaches for characterization of molecular drug resistance mechanisms, there have been recent reports of PZA resistant *M. tuberculosis* clinical isolates that encode wild type *pncA* and harbor mutations in other genes with possible roles in resistance^5–7^. An improved understanding of the molecular mechanisms that govern drug susceptibility and resistance will be crucial to advance genome-based molecular drug susceptibility testing and meet the challenge of ongoing trends in antimicrobial drug resistance.

Unlike other antimicrobial agents, PZA is only active against *M. tuberculosis* under specific environmental conditions *in vitro* and *in vivo*. Under standard culture conditions, PZA shows no growth inhibitory activity, whereas, exposure of bacilli to low pH is known to drive PZA susceptibility^8^. This observation is consistent with the finding that PZA shows no activity against *M. tuberculosis* within resting macrophages, yet, it is bactericidal against bacilli within activated macrophages that undergo greater phagosomal acidification^9^. Moreover, variable antitubercular PZA activity is observed in C3HeB/FeJ mice that show heterogeneous tuberculous lesions^10^. In these mice, PZA is efficacious in lesions showing acidic pH, but is ineffective against bacilli in lesions showing circumneutral pH^10^. Further, PZA shows sterilizing activity against *M. tuberculosis* in immune competent mice, yet, it lacks efficacy in athymic nude mice that are unable to mount a cell-mediated immune response^11^. Together, these studies demonstrate a pivotal role for host immunity in establishing sufficiently acidic microenvironments that are essential for the sterilizing antitubercular action of PZA^12^.

As a potential explanation for the pH-dependent action of PZA, it was proposed that POA might function as a proton ionophore^13^. By this model, PZA passively diffuses across the mycobacterial cell envelope and is then converted to POA anion by PncA. POA is then excluded from the cytoplasm by an unidentified efflux mechanism. A small proportion of POA (pKa 2.9) becomes protonated upon exposure to an acidic environment and reenters the cell by passive diffusion. Protons then dissociate from incoming molecules of POA and the cycle continues leading to dissipation of proton motive force and acidification of the cytoplasm. In support of this model, intrabacterial acidification and dissipation of the proton motive force have been observed following two days exposure of bacilli to PZA at pH 4.5^14^. However, it has not been demonstrated whether these events are due to POA acting as a protonophore or are the consequence of disruption of some other cellular process associated with maintenance of membrane potential. Importantly, proton motive force and intracellular pH are not measurably impacted by exposure of the bacilli to PZA at pH 5.8 which is typically used for PZA susceptibility testing^15^. Further, pH-independent PZA susceptibility can be achieved through overexpression of PncA^15^ or substitution of PZA with POA^15,16^. Thus, proton shuttling does not appear to be the basis for the pH-dependent action of PZA.

In addition to proton shuttling, several other modes of action for PZA have been proposed. These models include roles for POA in the inhibition of fatty acid synthesis, *trans*-translation, guanosine pentaphosphate synthetase/polyribonucleotide nucleotidyltransferase, quinolinic acid phosphoribosyltransferase and coenzyme A (CoA) biosynthesis ^17^. While these models are not mutually exclusive, direct inhibition of fatty acid biosynthesis and *trans*-translation by POA have been challenged by subsequent studies^18,19^. Multiple recent reports confirm a connection between PZA action and impairment of CoA synthesis in *M. tuberculosis*. Mutations in *panD*, encoding the first enzyme of the CoA biosynthesis pathway, have been shown to confer PZA and POA resistance^20–22^. Further, supplementation with intermediates of the CoA biosynthetic pathway was found to antagonize PZA and POA action^20,22,23^. Moreover, POA treatment has been shown to significantly decrease intracellular levels of CoA in *Mycobacterium bovis* strain BCG^22,24^. Based on ligand interaction and co-crystallography studies, it has been demonstrated that POA binds PanD and promotes its destabilization^25–27^. However, a *M. tuberculosis* strain deleted for *panD* shows measurable susceptibility to PZA^23^, indicating that PanD is not the only target of POA. Despite these recent advancements in the understanding of PZA mechanism of action, their connection to conditional PZA susceptibility has yet to be determined.

In this study, we used the genome-wide deep sequencing based approach, transposon sequencing (Tn-seq), to comprehensively interrogate which cellular pathways are associated with PZA susceptibility. Genetic associations were identified across various cellular processes, many of which have not been previously linked with PZA or POA resistance. Many of these functions play key roles in central metabolism and the cell envelope stress response. Further, we demonstrate that activation of this response through the extracytoplasmic function sigma factor E (SigE) is central to conditional PZA susceptibility and can be manipulated by exposing the bacilli to antibiotics that target peptidoglycan synthesis. These observations establish a paradigm shift in our understanding of the action of this important drug through defining the regulatory mechanism that underlies conditional PZA susceptibility.

## RESULTS

### Genome-wide analysis of molecular mechanisms for pyrazinamide resistance

Mutagenesis with the *himar1* transposon^28^ was used to investigate the genetic basis for mycobacterial PZA susceptibility. To avoid a preponderance of insertion mutations in *pncA* and circumvent the need for acidification of the growth medium, strains were selected for resistance to POA at pH 6.6. Similar conditions were recently used for the identification of spontaneous POA resistance mutations in *panD* and *clpC1* in *M. tuberculosis* and *M. bovis* BCG^22^. It is important to note that while POA shows inhibitory activity against *M. tuberculosis* at circumneutral pH, POA susceptibility is enhanced by exposure to low pH. Since we utilize the mycobacteriophage phAE180^29^ for delivery of *himar1*, and susceptibility to PZA and POA can be modulated by specific extracellular stimuli, the impact of phage infection on POA susceptibility was determined. On agar medium, the minimum inhibitory concentration (MIC) of POA for *M. tuberculosis* H37Rv was 200 μg ml^−1^ at pH 6.6 and 50 μg ml^−1^ at pH 5.8 (Table S1), consistent with previous reports^16,30^. Interestingly, when bacilli were infected with phAE180 and plated on solid medium (pH 6.6) containing kanamycin for selection of *himar1* transposon insertion, the POA MIC was 50 μg ml^−1^ (Table S1). Thus, like exposure to low pH, phage infection also enhances susceptibility of *M. tuberculosis* to this drug. Enhancement of POA susceptibility was also observed for *M. bovis* BCG following infection with phAE180 (Table S1), demonstrating that phage-mediated potentiation of mycobacterial POA susceptibility is not strain specific.

Spontaneous resistance to POA at circumneutral pH has been reported to occur at a frequency of 10^−5^ ^22^. When a pool of approximately 2×10^5^ independent *M. tuberculosis* H37Rv transposon insertion mutants was plated on 50 μg ml^−1^ POA at pH 6.6, 2×10^3^ colonies emerged, indicating that ~1% of insertions were associated with POA resistance. Fourteen independent isolates were chosen to assess POA and PZA resistance and to determine the corresponding chromosomal transposon insertion sites. From these isolates, 10 had unique insertions and 4 appeared to be siblings. Consistent with other recent studies^20–2,31,32^, insertions were identified in the carboxy-terminal coding region of *panD* and in the promoter region of *clpC1* (Table S2). Eight additional unique insertions were identified within seven other genes that had not previously been associated with PZA or POA resistance (Table S2). These strains showed 2-to 16-fold resistance to POA at pH 6.6 (MIC 400 to 3200 μg ml^−1^), whereas PZA resistance was typically only 2-fold at pH 5.8 (MIC 100 μg ml^−1^) relative to the parental strain (Table S2), as was previously described for *panD* and *clpC1* mutant strains^31^. It is noteworthy that despite ongoing debate over the breakpoint concentration for PZA, resistance of *M. tuberculosis* clinical isolates to >50 μg ml^−1^ is associated with poor sputum conversion rates^33^. For all mutant strains that were tested, INH susceptibility was indistinguishable from that of the parental strain (Table S2), indicating that resistance was specific to PZA and POA.

To comprehensively identify genes associated with POA susceptibility, independently prepared pools of saturated *M. tuberculosis* H37Rv transposon insertion mutants were plated as described above without or with 50 μg ml^−1^ POA in duplicate. In the absence of POA, approximately 2×10^5^ colonies were recovered from each pool. Similar to that described above, selection with POA yielded approximately 2×10^3^ POA resistant colonies from each pool. To identify transposon insertion sites in these libraries, colonies were collected, genomic DNA was extracted, transposon adjacent regions were enriched by PCR and deep sequencing was performed^34^. Sequencing of non-POA selected libraries yielded 1.1 M and 0.72 M high quality reads that mapped to 46,901 and 34,635 unique TA sites, respectively (Fig. 1a, Dataset S1). Sequencing of POA selected libraries resulted in 0.97 M and 0.48 M high quality reads that mapped to 9,903 and 1,058 unique TA sites, respectively (Dataset S1). Insertion sites that were present at an abundance of ≥5 read counts in both POA-selected libraries were analyzed further. These insertions constituted 93% of total reads and mapped to 275 TA sites within 58 genes and intergenic regions (Fig. 1b, Table S3). The remaining 7% of reads were low abundance (<5 read counts, 1% of reads) and/or only observed in one replicate (6% of reads), and were excluded from further analysis. Fold enrichment for insertions in highly represented loci was determined by comparing the mean relative read abundance to that from the no POA condition (Fig. 1c, Table S3). Twenty genes showed a level of enrichment between 2 and 1000-fold in the presence of POA and achieved a threshold of significance of <0.05 (Fig. 1c, Table S3).

**Figure 1.**
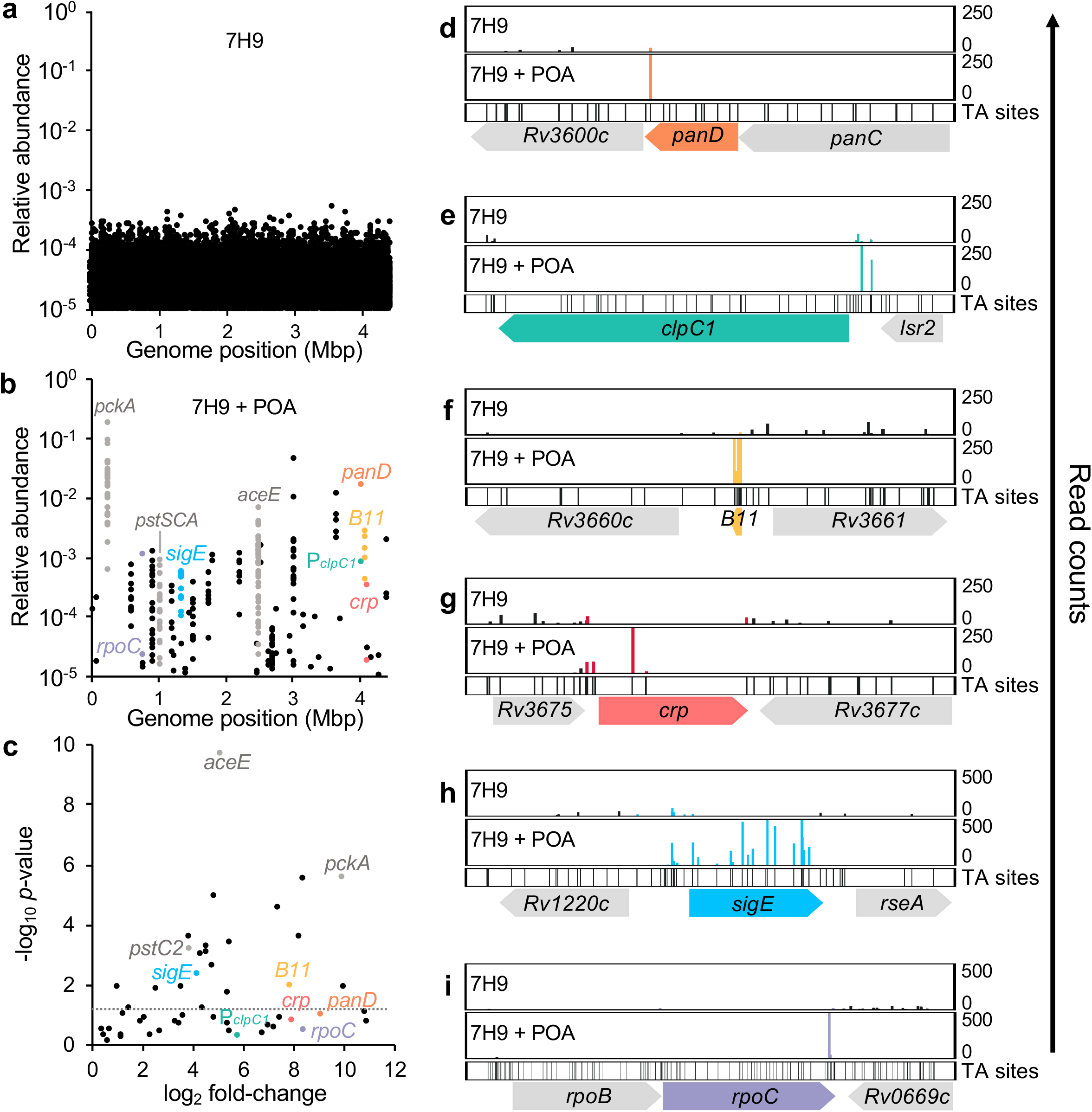
Genes associated with *M. tuberculosis* PZA susceptibility by Tn-seq. Libraries of 2×10^5^ independent *M. tuberculosis* H37Rv *himar1* insertion mutants (4-fold saturation) were plated on 7H9 agar without (**a**) or with POA (**b**). Genomic DNA was extracted, processed and sequenced as described in Minato, *et al* 2019 *mSystems*^34^. **a** & **b** show mean abundance relative to total TA insertion read counts from two independent replicates. Panel **c** shows log_2_ fold change in mean relative abundance of **b** compared to **a** by gene and respective-log_10_ *p* value (dotted line at *p* = 0.05). Read count comparisons for *panD* (**d**), P_*clpC1*_ (**e**), *B11* (**f**), *crp* (**g**), *sigE* (**h**) and *rpoC* (**i**) are shown.

The majority of highly represented insertions were identified in loci that are functionally associated with intermediary metabolism and respiration, cell wall and cell processes, information pathways, stable RNAs, virulence-detoxification-adaptation, and conserved hypothetical proteins (Table 1). As anticipated, highly abundant insertions were identified in the 3’ end of *panD* (Fig. 1b,c,d) and promoter of *clpC1* (Fig. 1b,c,e), yet, these insertions fell below the significance threshold due to a limited number of available TA sites. Numerous insertions in genes for the persistence-associated high affinity phosphate transport system^35,36^ were observed (Fig. 1b,c, Fig. S1a), consistent with reports of point mutations in *pstC2* in PZA resistant laboratory and clinical isolates^6,32^. Further, highly abundant insertion sites also included eight of the ten genes that were identified in our primary analysis described in Table S2. More than 80% of highly represented insertions in this analysis were located in genes involved in central carbon metabolism. Insertions throughout *pckA* (encoding phosphoenolpyruvate carboxykinase, the first enzyme of gluconeogenesis) constituted the vast majority (73%) of reads (Fig. 1b,c, Fig. S1b). The second most abundant set of insertions were found throughout *aceE* (encoding the E1 subunit of pyruvate dehydrogenase) and represented 5.3% of all reads (Fig. 1b,c, Fig. S1c). To confirm that loss-of-function mutations in *pckA* and *aceE* can confer POA resistance, these genes were deleted from H37Rv using the recombineering-based approach ORBIT^37^. Both strains were found to be at least 4-fold resistant to POA (MIC ≥800 μg ml^−1^) relative to H37Rv (MIC 200 μg ml^−1^). Less abundant insertions were reproducibly observed in the promoter region for *dlaT* (encoding the E2 subunit of pyruvate dehydrogenase), and within *icd2* (encoding isocitrate dehydrogenase) and *kgd* (encoding alpha-ketoglutarate decarboxylase) (Table 1, Table S3) indicating an important association between POA action and central carbon metabolism. Growth at low pH is known to result in extensive metabolic remodeling that is dependent upon PckA^38^, which may be an important event linked to PZA potentiation. Thus, it is likely that these mutations mitigate the deleterious impact of POA on CoA biosynthesis through increasing availability of oxaloacetate for production of L-aspartate, the precursor of β-alanine. Consistent with this model, we find abundant insertions in *B11* (*ncRv13660c*; Fig. 1b,c,f) a *trans*-encoded small RNA that regulates CoA synthesis through duplex formation with *panD* mRNA^39^. Insertions were also observed in *crp* (*Rv3676*; Fig. 1b,c,g) encoding the transcriptional regulator of *B11*^40^. In addition, insertions were also identified in genes for the sulfate transport system encoded by the *subI-cysTWA1* operon (Table 1, Fig. S1d). In contrast to our findings with POA, this operon was recently described for its role in intrinsic tolerance to many other antitubercular drugs^41^. Similarly, *mmpS3* has also been implicated in broad drug tolerance of *M. tuberculosis*^41^, whereas we find its inactivation to be associated with POA resistance (Table S3). It is intriguing that *mmpS3* plays a key role in modulation of cell wall synthesis, as abundant insertions were found in *sigE* (Fig. 1b,c,h), encoding the cell envelope stress response sigma factor, and in the carboxy-terminal domain of *rpoC* (Fig. 1b,c,i) which mediates critical interactions between SigE and RNA polymerase^42^. Together, these observations indicate contrasting roles for genes associated with POA susceptibility versus tolerance to other drugs that target actively replicating bacilli, consistent with the long-standing notion that PZA predominantly targets persister populations.

**Table 1.**
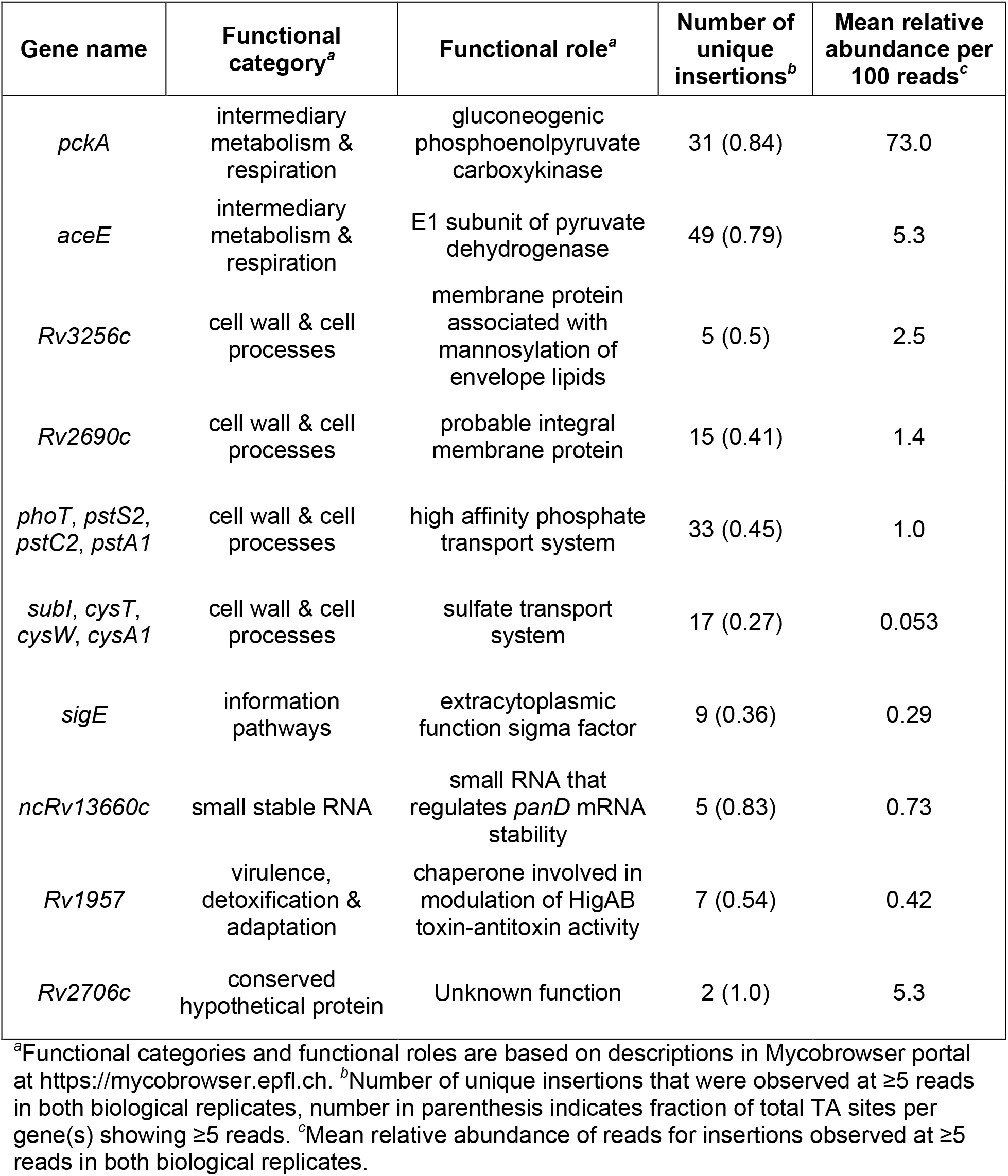
Loci of highly abundant transposon insertions associated with pyrazinoic acid resistance in *M. tuberculosis*

### Activation of the cell envelope stress response potentiates PZA susceptibility

Based on the new findings described above and the established role of SigE in response to low pH ^43^, it is likely that acidic conditions drive PZA and POA susceptibility through activation of the cell envelope stress response. To confirm a role for this response as a driver of PZA and POA action, susceptibility of a strain deleted for *sigE* (Δ*sigE*) was tested in liquid medium. For wild type *M. tuberculosis* H37Rv the MIC of PZA was 50 μg ml^−1^ and POA was 200 μg ml^−1^ (Table 2). In contrast, the MIC of PZA was 400 μg ml^−1^ and POA was ≥800 μg ml^−1^ for the *M. tuberculosis* Δ*sigE* strain (Table 2), confirming SigE as a critical driver of susceptibility. Expression of *sigE* from an integrating vector was sufficient to restore susceptibility of the Δ*sigE* strain to both PZA and POA (Table 2). INH susceptibility was indistinguishable for all strains indicating that the association between *sigE* and drug resistance is specific to PZA and POA (Table 2). Indeed, in contrast to the role for SigE in conditional susceptibility to PZA, recent work by Pisu and colleagues demonstrates that the SigE response is important for mediating tolerance of *M. tuberculosis* to numerous other antimicrobial agents ^44^.

**Table 2.** The SigE response plays a central role in PZA and POA susceptibility of *M. tuberculosis*.

| *M. tuberculosis* strain | Relevant characteristic | POA MIC<sup>a</sup> (µg ml<sup>-1</sup>) | PZA MIC (µg ml<sup>-1</sup>) | INH MIC (µg ml<sup>-1</sup>) |
| --- | --- | --- | --- | --- |
| H37Rv | Parental strain | 200 | 50 | 0.0625 |
| H37Rv $\Delta sigE$ | H37Rv deleted for sigma factor E gene | ≥800 | 400 | 0.0625 |
| H37Rv $\Delta sigE$ pMV306-*sigE* | $\Delta sigE$ strain expressing *sigE* in trans | 400 | 50-100 | 0.0625 |

<sup>a</sup> Minimum inhibitory concentration (MIC) is defined as the minimum concentration required to inhibit 90% of growth relative to the no drug control after 14 days (for POA and PZA) or 7 days (for INH) of incubation at 37 °C. POA and INH exposure was performed at pH 6.6, PZA exposure was performed at pH 5.8.

Under non-stress conditions, the cell envelope stress response is muted through sequestration of SigE by the anti-sigma factor RseA ^45^. Upon sensing of cell envelope stress, RseA is phosphorylated by the essential serine-threonine kinase PknB and is subsequently degraded by the ClpC1P2 protease resulting in release of SigE thereby promoting expression of its regulon ^46^. To further evaluate the role of the SigE response in susceptibility of *M. tuberculosis* to PZA and POA, a strain deleted for *rseA* (Δ*rseA*) was assessed for PZA and POA susceptibility under acidic (pH 5.8) and circumneutral (pH 6.6) incubation conditions. As anticipated, under acidic conditions, the Δ*rseA* strain showed a level of PZA and POA susceptibility that was indistinguishable from the wild type control (Fig. 2a,b). Also, as has been previously described, wild type H37Rv was exposed to concentrations up to 800 μg ml^−1^ PZA at circumneutral pH, growth was not notably impaired (Fig. 2c). In striking contrast, when the Δ*rseA* strain exposed to PZA under circumneutral conditions (Fig. 2c), it showed a level of susceptibility comparable to that which was observed under acidic conditions (Fig. 2a). Similarly, susceptibility of the Δ*rseA* strain to POA was equivalent under circumneutral (Fig. 2d) and acidic conditions (Fig. 2b). Importantly, INH susceptibility was unchanged (Fig 2e), indicating that the potentiation effect through constitutive activation of the SigE response is specific to PZA and POA. Taken together, these results demonstrate that activation of the SigE cell envelope stress response is both necessary and sufficient for driving PZA and POA susceptibility in *M. tuberculosis*.

**Figure 2.**
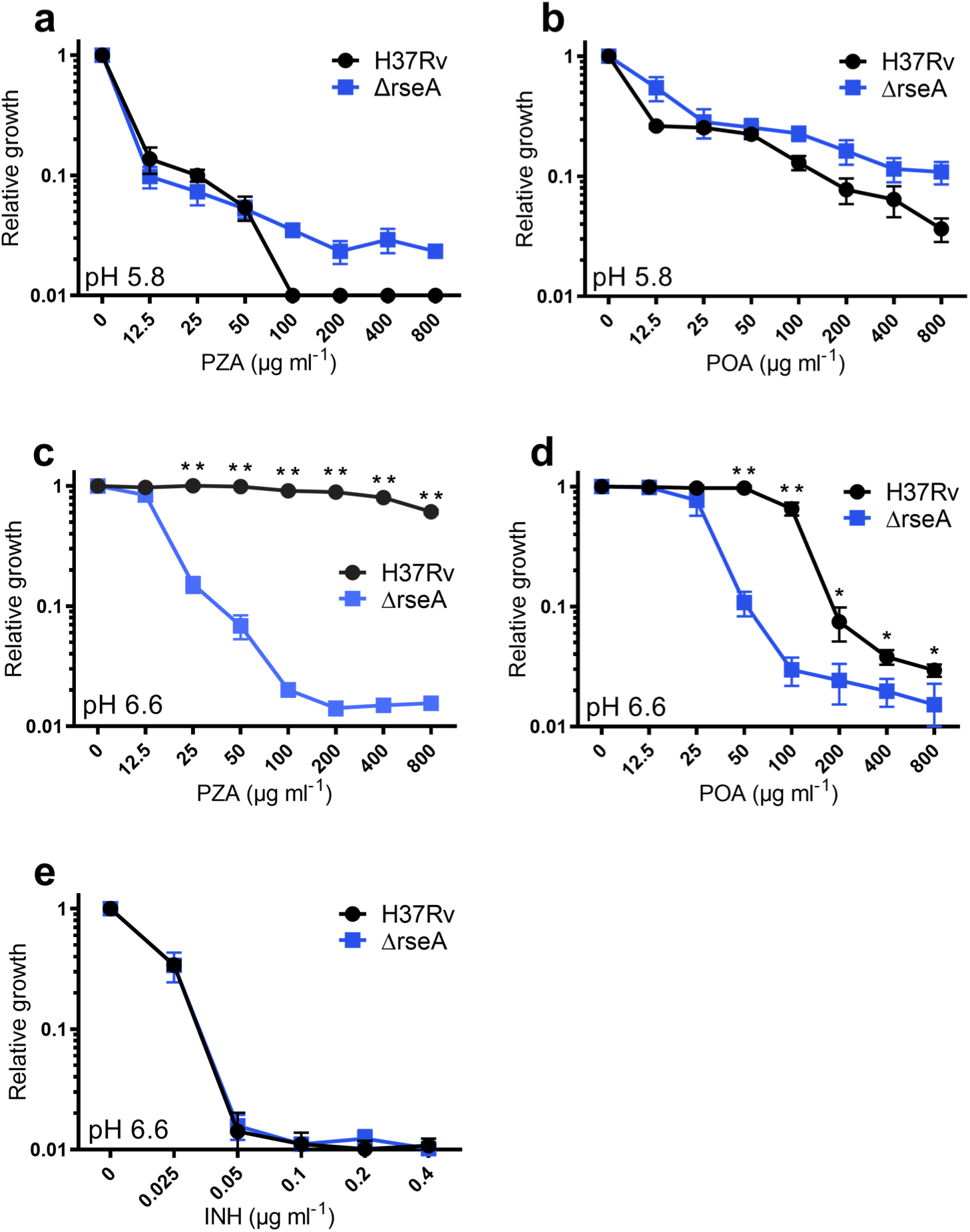
Deletion of *rseA* in *M. tuberculosis* confers constitutive susceptibility to PZA. *M. tuberculosis* H37Rv (wild type, black circles) and the Δ*rseA* strain (blue) were grown in 7H9 at pH 5.8 (**a, b**) or pH 6.6 (**c, d, e**) and exposed to PZA (**a, c**) or POA (**b**, **d**) for 14 days or INH for 7 days (**e**). OD_600_ was measured, relative growth was determined by dividing by the value of the no drug control. Plotted values represent the mean ± S.D. of 3 biological replicates. \**p* <0.05, \*\**p* <0.0002 by 2 tailed Student’s *t*-test.

Next, we evaluated whether there were any signatures of selection for *sigE* mutations in PZA-resistant clinical isolates. Considering that most clinical PZA resistance is mediated by *pncA* mutations and an intact SigE regulon is essential for tolerating other antitubercular therapeutics ^44^, we reasoned that any selection for *sigE* mutants would be exceedingly rare and appear more frequently—if not exclusively—in *pncA_WT_* PZA monoresistant isolates. Accordingly, we contrasted the prevalence of *sigE* mutations in a large global set of clinical isolates (n=1,215) to a recently curated set of PZA mono-resistant isolates (n=18) in which *pncA_WT_* isolates were markedly overrepresented ^47^ including, three that are both *pncA_WT_* and *panD_WT_*. Among all isolates, six non-synonymous *sigE* mutations were observed. Among these, *sigE*_V166L_ was most tenable for conferring an advantageous functional effect. The V166L substitution falls within the SigE σ_2_/σ_4_ linker (Fig. 4), which was recently shown to interface with template single-stranded DNA in the active center cleft of RNA polymerase, and influences open complex formation, abortive production and promoter escape during transcription initiation ^48^. A mutation in in this linker domain could affect how SigE associates with promoters of its regulon, ostensibly altering the conditionality or specificity of its transcription-activating action without uniformly deleterious effects. The single isolate harboring this mutation was one of only three PZA-monoresistant, PncA_WT_, and PanD_WT_ isolates, a statistically improbable observation if the mutation and atypical basis of PZA resistance in the isolate harboring it were independent (*p* = 0.014, Fisher’s Exact corrected for multiple hypotheses). The presence of this mutation in a lone isolate monoresistant to PZA and lacking the two most established resistance-conferring clinical mutations suggests a potential role in PZA resistance. This finding is consistent with the idea that the SigE-dependence of PZA susceptibility we have characterized *in vitro* also operates within the context of human infection.

### Peptidoglycan targeting agents strongly potentiate PZA antitubercular action

Based on the observation that the SigE response governs PZA conditional susceptibility, we reasoned that specific activation of this response through co-treatment with cell envelope damaging agents should lead to potentiation of PZA activity against *M. tuberculosis*. To assess the impact of differing types of cell envelope stress on PZA activity, compounds disrupting synthesis of various layers within the mycobacterial cell envelope were evaluated for fractional inhibitory concentration index (FICI) values using checkerboard assays in standard 7H9 medium at pH 6.6. Meropenem and D-cycloserine were chosen to target peptidoglycan biosynthesis. Meropenem, a β-lactam, irreversibly inhibits penicillin binding proteins (PBPs) thereby preventing cross linking of the peptidoglycan side-chains. The β-lactamase inhibitor clavulanate was included to inhibit the *M. tuberculosis* β-lactamase and improve stability of meropenem^49^. Combining meropenem/clavulanate with PZA in a checkerboard assay yielded synergistic FICI values of 0.265 and 0.5 for *M. tuberculosis* H37Rv (Fig. 3a) and *M. tuberculosis* Erdman (Fig. 3b), respectively. Further, the minimum FICI value achieved for the H37Rv Δ*sigE* was 0.875, indicating no drug interaction (Fig. 3c). Thus, the synergistic activity between meropenem and PZA is dependent upon the SigE response.

**Figure 3.**
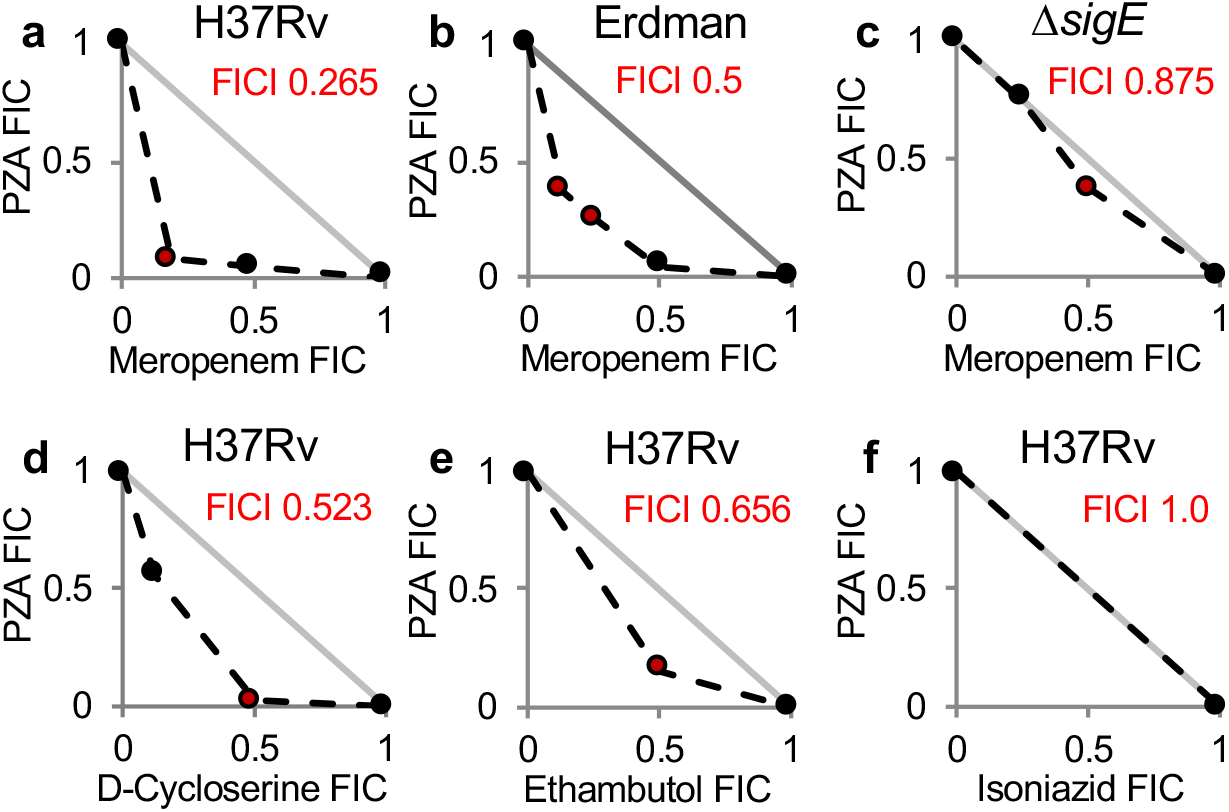
Peptidoglycan synthesis inhibitors potentiate antitubercular activity of pyrazinamide. *M. tuberculosis* H37Rv (**a, d-f**), Erdman (**b**) and H37Rv Δ*sigE* (**c**) were grown in 7H9 at pH 6.6 with varying concentrations of PZA and either, meropenem/clavulanate (**a,b,c**), D-cycloserine (**d**), ethambutol (**e**), or isoniazid (**f**) in checkerboard format. OD_600_ was determined after 7 days of incubation. Plots were generated based on the average fractional inhibitory concentration (FIC) calculated from two biological replicates. The lowest FIC index (FICI) values are indicated in red. Line of additivity is shown in grey (FICI of 1).

Based on the observed synergy between meropenem and PZA, we assessed whether the mechanism of peptidoglycan damage impacted potentiation of PZA action. D-cycloserine is an amino acid analogue that targets L-alanine racemase and D-alanyl-alanine synthetase, preventing synthesis of new peptidoglycan, as opposed to blocking PBP crosslinking. Checkerboard assays for D-cycloserine and PZA yielded a minimum FICI value of 0.523 (Fig. 3d). While not meeting the threshold for synergy, these results indicate a strong, unidirectional potentiation of PZA susceptibility by D-cycloserine.

Next, we interrogated whether drugs targeting other mycobacterial envelope layers were also capable of enhancing susceptibility to PZA. EMB was chosen to target the arabinogalactan layer. EMB targets the arabinosyl transferases which is essential for synthesis of the arabinogalactan layer of the mycobacterial envelope ^50^. Co-treatment with EMB also did not potentiate PZA activity (Fig. 3e). The lowest FICI value achieved was 0.656, indicating drug additivity. INH was chosen to target the mycolic acid layer. INH inhibits InhA, the enoyl acyl carrier protein (ACP) reductase, preventing mycolic acid synthesis^51^. Co-treatment with INH did not have a measurable impact on PZA susceptibility (Fig. 3f). These observations are in agreement with previous observations that INH and PZA do not interact ^52^.

## DISCUSSION

Despite several decades of study and clinical use, the mechanistic basis for conditional susceptibility of *M. tuberculosis* to PZA remained elusive. In this study, we utilized a genome-scale approach to reveal cellular responses that drive PZA susceptibility and are orchestrated through activation of the cell envelope stress response that is governed by the alternate sigma factor SigE (Figure 5). Indeed, we demonstrate that constitutive activation of this stress response through deletion of the anti-sigma factor gene *rseA* renders *M. tuberculosis* intrinsically susceptible to PZA, whereas, blocking this response leads to resistance. Resistance was also observed when activation of the SigE response was impaired through disruption of the *clpC1* locus which plays a central role in degradation of RseA. Moreover, we demonstrated that activation of the cell envelope stress response through co-treatment with peptidoglycan damaging agents leads to potent potentiation of PZA and was SigE-dependent. These observations offer a stark contrast to the relationship between the SigE response and other antimicrobial agents where this response has been shown to promote tolerance to numerous other antitubercular drugs such as INH, EMB, RIF, and streptomycin^44^. The precise role that SigE has in promoting survival during stress treatment has not been fully elucidated, but recent studies have suggested that the regulon may play a role in maintenance of *M. tuberculosis* dormancy^53^. SigE-dependent maintenance of dormancy may be accomplished through modulation of central metabolism^54^ or through the MprAB-SigE driven regulation of a Psp-like system that maintains *M. tuberculosis* cell envelope integrity^55^. These connections between SigE and dormancy that lead to increased tolerance to multiple antibiotics may be a double-edged sword in the case of PZA treatment, offering a potential explanation for PZA efficacy against *M. tuberculosis* independent of growth state.

**Figure 4.**
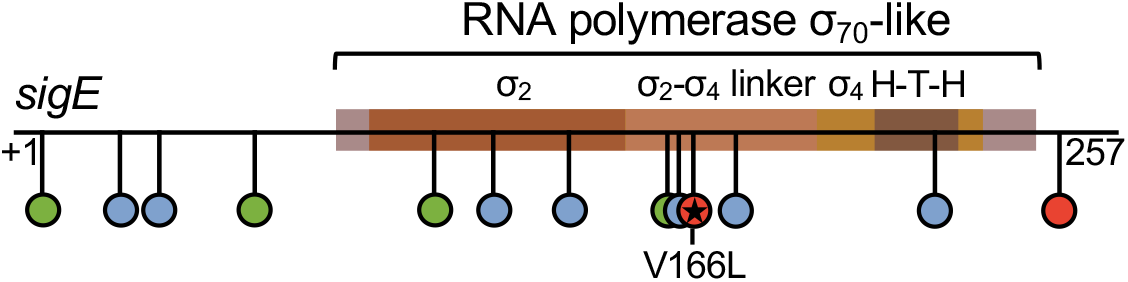
Distribution of Tn-seq mutants and natural sequence polymorphisms observed in clinical isolates in SigE functional domains. Location of Tn-seq insertions enriched under POA pressure non-synonymous SNPs observed across in PZA mono-resistant clinical isolates with respect to SigE domain architecture imported from InterPro and curated from recent literature ^48^. Locations of all missense mutations present in at least one isolate and all TA insertion sites with greater relative abundance in POA-containing media than 7H9 are depicted.

**Figure 5.**
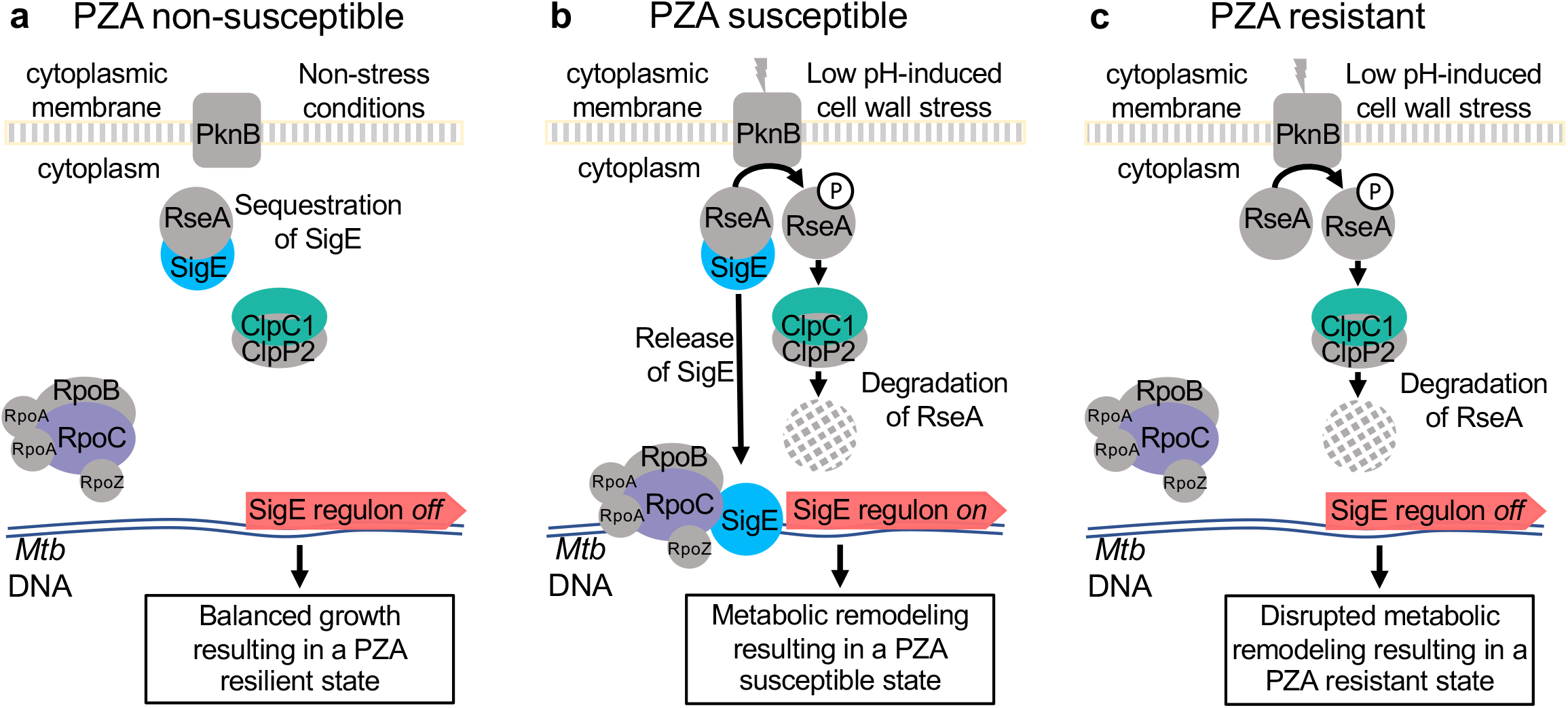
Model for low pH-mediated potentiation of PZA susceptibility. (**a**) Under conditions of balanced growth, *M. tuberculosis* is in a non-stressed state resulting in PZA resilience. (**b**) Under conditions of low pH or other cell wall stress inducing conditions, signaling through PknB mediates phosphorylation of RseA which is subsequently degraded by ClpC1P2 liberating SigE. Expression of the SigE regulon results in metabolic remodeling that poises bacilli in a PZA susceptible state. (**c**) Failure to activate the SigE response prevents cell wall stress driven metabolic remodeling resulting in PZA resistance.

One counterintuitive finding in our study is the observation that disruption of ClpC1 and SigE conferred measurable resistance to POA in the absence of external cellular stress. This finding is not unprecedented given the well-established role of PknB in activation of the SigE response through phosphorylation of RseA^46^. PknB is an essential transmembrane serine/threonine-protein kinase that contains an extracellular peptidoglycan sensing domain^56^. Through sensing peptidoglycan fragments present at the septum, PknB plays a direct role in determining cell shape and controlling cell division^57–59^. It stands to reason that the SigE response might be transiently activated during cell division in addition to its well-described role in stress response. Activation of the SigE regulon during cell division could explain *M. tuberculosis* susceptibility to POA under non-stress conditions, as well as cases where pH has been uncoupled from PZA and POA susceptibility^15^.

Currently, the biochemical event linking activation of the SigE regulon to PZA susceptibility is unclear. Many of the highly abundant hits in our screen were mapped to genes capable of impacting cellular CoA levels. These observations are consistent with previous reports linking mutations in CoA biosynthetic genes as well as supplementation with CoA precursor molecules to PZA resistance^20,22,23^. Gopal and colleagues have recently shown that intracellular CoA pools play a major role in determining PZA and POA susceptibility^22^. Phosphoenolpyruvate carboxykinase, α-ketoglutarate dehydrogenase and pyruvate dehydrogenase play key roles in central carbon metabolism where CoA is used at various steps to move carbon into, through and out of the TCA cycle. Changes in the proteome during stress response conditions are expected to impact CoA-dependent metabolism. Exposure to acidic pH leads to specific metabolic remodeling that provides a fitness advantage^38^. The SigE response likely orchestrates this adaptation through major transcriptional remodeling^44,60^. These data, taken with the previous finding that POA treatment decreases intracellular abundance of CoA^22^, suggest that perturbations in CoA levels influence *M. tuberculosis* susceptibility to PZA and POA. We propose that cell envelope stress leads to an induction of the SigE regulon, which in turn modulates cellular CoA levels. Once CoA levels have been reduced, treatment with PZA or POA further depletes CoA, leading to growth inhibition through metabolic dysfunction (Figure 5). Further support for this model is the observation that supplementation with exogenous CoA precursors can antagonize PZA and POA action^20,22,23^. These observations also suggest that CoA-dependent metabolism may determine the impact of PZA and POA treatment. Under conditions that do not induce the SigE response, PZA and POA are needed at relatively high concentrations to exert a strong enough effect on intracellular CoA and impact growth. Yet, under conditions that trigger the SigE response, intracellular CoA levels are more vulnerable to perturbations by PZA or POA.

The ability to disconnect PZA susceptibility from pH allows for isolation of informative resistant mutants that have been missed by previous screens and selections. There may be an *in vivo* fitness cost providing a strong counter selection against certain mutations during infection. For example, both *panD* and *aceE* have been characterized as essential genes for growth in a mammalian host^61,62^. These findings offer new insight into characterization of clinically resistant strains that do not have mutations in loci that are commonly associated with PZA resistance, such as *pncA*. A recently sequenced PZA clinical isolate described by Maslov *et al.* harbored 15 non-synonymous mutations in protein coding sequences^5^, and it was concluded that further functional analysis would be needed to associate any of these mutations to the observed PZA resistance. Combined with our findings, it is possible to correlate these putative resistance mutations with several highly abundant loci detected in our study, namely *aceE, rpoC,* and *dlaT* as candidates contributing to PZA resistance. Independently, each of these mutations is likely to confer a low level of PZA resistance. However, in combination, the effect of these mutations may be additive and culminate in a high level of resistance to PZA. Whole genome based polygenic surveys for loci associated with PZA resistance may offer a powerful predictive tool for rapid molecular based PZA susceptibility testing.

This new understanding of conditional PZA susceptibility may be particularly impactful in the context of TB therapy in individuals with compromised immunity. During TB infection, bacilli are engulfed by dendritic cells and alveolar macrophages where they reside within phagosomal compartments^63^. Prior to activation by helper T cell signals, the bacilli are able to replicate within phagosomes despite conditions of mild acidity (pH 6.2 − 6.4) and exposure to sub-inhibitory levels of reactive oxygen intermediates^63^. Following cell-mediated activation by pro-inflammatory cytokines, such as interferon-γ and tumor necrosis factor-α, intracellular bacilli must adapt to more severe growth-inhibitory stressors, such as lowered pH (pH 4.5 − 5.4), nutrient limitation and bombardment with high levels of reactive oxygen and reactive nitrogen intermediates^63^. While T cell-mediated activation of macrophages is not usually sufficient to eliminate all bacilli from the host, it is essential for long-term containment in the form of a latent infection^63^. Impairment of CD4^+^ T cell signals through prolonged antigen stimulation, genetic lesions in the interferon-γ signaling pathway, anti-tumor necrosis factor-α therapy and HIV co-infection are common mechanisms that drive progression to overt TB disease and other mycobacterial infections^64^. Consistent with a role for host involvement in the sterilizing activity of PZA, it has recently been demonstrated that PZA-mediated killing is impaired in athymic nude mice that are unable to drive cell-mediated activation of monocytes due to a complete lack of T cells^11^. Combined with our findings, this essential role for T cell responses for *in vivo* PZA action suggests that activation of host antimicrobial stressors are critical for driving PZA susceptibility of *M. tuberculosis*. As such, these observations indicate that PZA might have suboptimal activity in individuals with compromised cell-mediated immunity. Our findings open up potential avenues to improve PZA efficacy in the context of immune deficiency through the adjunctive use of carbepenems to drive the cell envelope stress response.

## METHODS

### Bacterial strains and growth conditions

Middlebrook 7H9 medium (Difco) supplemented with 10% (vol/vol) oleic acid-albumin-dextrose-catalase (OADC, Difco), 0.2% (vol/vol) glycerol, and 0.05% (vol/vol) tyloxapol (Sigma) or 7H10 medium (Difco) supplemented with 10% (vol/vol) OADC and 0.2% (vol/vol) glycerol were used to cultivate *M. tuberculosis* H37Rv and Erdman and *M. bovis* BCG. When necessary, kanamycin (Thermo Scientific) and/or hygromycin (Corning) were added at 50 and 150 μg ml^−1^, respectively.

### Transposon mutagenesis and transposon sequencing

*M. tuberculosis* H37Rv was mutagenized with the mariner *himar1* transposon using the temperature-sensitive mycobacteriophage phAE180^29^. Approximately 2×10^5^ independent transposon mutagenized bacilli were spread on Middlebrook 7H9 medium supplemented with 10% (vol/vol) OADC and 0.2% (vol/vol) glycerol containing 1.5% agar (Difco), 50 μg ml^−1^ kanamycin, and without or with 50 μg ml^−1^ POA (Sigma) in 245 mm^2^ BioAssay Dishes (Nunc). For analysis of individual POA resistant strains, secondary plating was performed on supplemented 7H10 medium without or with 400 μg ml^−1^ POA.

For transposon sequencing, plates were incubated for two weeks, colonies were collected and genomic DNA was extracted as previously described^34^. Briefly, colonies were scraped into 10 ml of 7H9 complete medium, the mycobacteria were pelleted, and incubated at 80 °C for 2 hours. Cultures were pelleted, re-suspended and washed in 500 μl of 25 mM Tris pH 7.9, 10 mM EDTA, and 50 mM glucose. The cells were re-suspended in 450 μl of 25 mM Tris pH 7.9, 10 mM EDTA, and 50 mM glucose with 50 μl of 10 mg ml^−1^ lysozyme. Samples were incubated at 37 °C for 16 hours. Next, 100 μl of 10% SDS and 50 μl of 10 mg ml^−1^ proteinase K were added and the mixture was incubated at 55 °C for 30 min. 200 μl of 5 M NaCl and 160 μl of CTAB saline solution (0.7 M NaCl, 0.275 M hexadecyl-trimethylammonium bromide, CTAB) were added and the samples incubated at 65 °C for 10 min. DNA was extracted using multiple chloroform:isoamyl alcohol treatments. The DNA was precipitated with isopropanol and washed with 70% ethanol prior to its resuspension in EB buffer (Qiagen). DNA fragmentation and Illumina P7 adaptor (CAAGCAGAAGACGGCATACGAGAT) ligation were performed in NeoPrep Library Prep System (Illumina). Transposon junctions were amplified by using a transposon specific primer Mariner_1R_TnSeq_noMm (TCGTCGGCAGCGTCAGATGTGTATAAGAGACAGCCGGGGACTTATCAGCCAACC) and P7 (CAAGCAGAAGACGGCATACGAGAT) primers with HotStarTaq Master Mix Kit (Qiagen). The following PCR condition was used; 94 °C for 3 min, 19 cycles of 94 °C for 30 seconds, 65 °C for 30 seconds, and 72 °C for 30 seconds, 72 °C for 10 minutes. The *himar1* enriched samples were diluted 1:50 and amplified by using a P5 indexing primer (AATGATACGGCGACCACCGAGATCTACAC[i5]TCGTCGGCAGCGTC, [i5] barcode sequence) and P7 primer HotStarTaq Master Mix Kit (Qiagen) to add unique barcodes and the necessary P5 and P7 flow cell adaptor sites for Illumina sequencing. The following PCR conditions were used; 94 °C for 3 min, 94 °C for 30 seconds, 55 °C for 30 seconds, 72 °C for 30 seconds. Sequencing was performed on Illumina MiSeq by the University of Minnesota Genomics Center. Transposon and adapter sequences were trimmed from the 5’ end of sequencing reads using CutAdapt ^65^. We also discarded all the sequence reads that did not contain adaptor sequence in the 5’ trimming process. After the 5’ trimming process, all the sequence reads begin with TA. Adapter sequences were trimmed and sequence reads that were shorter than 18 bp were discarded. The default error rate of 0.1 was used for all trimming processes. The trimmed sequence reads were mapped (allowing 1 bp mismatch) to the *M. tuberculosis* H37Rv genome (GenBank: AL123456.3) using Bowtie^66^. The genome mapped sequence reads were printed in SAM format and counted sequence reads per each TA dinucleotides site in the *M. tuberculosis* H37Rv genome using SAMreader_TA script^67^.

### Determination of drug susceptibility

Strains were cultured to mid-log phase and subsequently inoculated at an initial OD_600_ of 0.01 into supplemented 7H9 medium at the indicated pH. The minimum inhibitory concentration (MIC) was defined as the minimum concentration of drug required to inhibit at least 90% of growth relative to the drug free controls, and was determined by measuring OD_600_. Drug susceptibility testing for PZA and POA was carried out using media at pH 5.8 or pH 6.6 with 14 days of incubation at 37 °C. INH MIC determinations were carried out in medium at pH 6.6 with 10 days of incubation at 37 °C. POA agar MIC determinations were conducted using 7H10 medium at pH 6.6. *M. tuberculosis* strains were spot diluted onto plates containing various concentrations of POA. MIC was determined at day 14 by assessing colony growth.

### Determining mycobacteriophage-mediated potentiation of POA activity

To determine the impact of mycobacteriophage transduction on POA susceptibility, *M. tuberculosis* and *M. bovis* strains were grown to OD_600_ 0.5. Cultures were pelleted by centrifugation and washed twice in an equal volume MP buffer (50 mM Tris, 150 mM NaCl, 10 mM MgCl_2_, 2 mM CaCl_2_). Cell pellets were re-suspended in 1 ml of MP buffer containing phage at ≥10^10^ plaque forming units ml^−1^ for a multiplicity of infection of 10. The cell/phage re-suspension was incubated for 24 hours at 37 °C in atmospheric CO_2_. After 24h, cells were re-suspended by gently pipetting 10 times and serially diluted. Dilutions were plated on supplemented 7H10 agar plates containing kanamycin and varying concentrations of POA. Colonies were counted at day 14 to assess POA susceptibility. A kanamycin resistant empty vector strain (*M. tuberculosis* strain H37Rv pUMN002) was treated identically with MP buffer containing no phage for comparison as a vehicle control.

### Construction of *M. tuberculosis* H37Rv deletion mutant strains

Deletion of *aceE*, *pckA* and *rseA* in *M. tuberculosis* H37Rv was accomplished using the ORBIT recombineering system^37^. In brief, *M. tuberculosis* H37Rv cells previously transformed with pKM461, encoding tetracycline-inducible Che9c RecT annealase and Bxb1 integrase, were grown to mid-log phase (OD_600_ ~ 0.8) in supplemented 7H9 medium containing 50 μg ml^−1^ kanamycin for selection of pKM461 containing cells. Once an OD of ~0.8 was reached, anhydrotetracycline was added to a final concentration of 500 ng ml^−1^ and allowed to incubate for 8 hours. Following induction, 2M glycine was added to the culture and allowed to incubate further overnight (~16 hr) while shaken at 37 °C. The next day, cells centrifuged in 50 ml conical tubes at 4,300 rpm for 10 minutes to pellet cells. Cell pellets were resuspended in an equal volume of 10% glycerol. The centrifugation and washing steps were repeated. A final centrifugation and washing step were performed with cells being resuspended in 3 ml of 10% glycerol. Cells were electroporated with 1 μg of a targeting oligonucleotide containing an *attP* site core and 200 ng of the knockout plasmid pKM464 carrying an *attB* site and hygromycin resistance marker. The targeting oligonucleotide sequence for deletion of *aceE* was CCCGACCGAGTTCGGGTGATCCGCGAGGGTGTGGCGTCGTATTTGCCCGACATTGATCCC<u>GGTTTGTCTGGTCAACCACCGCGGTCTCAGTGGTGTACGGTACAAACC</u>CAGACCACGGATCCCGGTCCCGGGGCCTAACGCCGGCGAGCCGACCGCCTTTGGCCGAAT, for deletion of *pckA* was TGCGTGCGGGGGCTTATGCGTCTGCTCGCCCTAACCTAGGCGCTCCTTCAGGGCGTCGAA<u>GGTTTGTACCGTACACCACTGAGACCGCGGTGGTTGACCAGACAAACC</u>ATCCAGACCGGGGATGGTCGCTGAGGTCATCGAATTCTCCTGCGTAGTTATCGGGTGCTC and for deletion of *rseA* was GCAGCAACCCCGCCATGCGCTGCGACAAGTGGCTCACCGGCTAGCGACGCACCCGCGATTGCCGGCCCCG<u>GGTTTGTACCGTACACCACTGAGACCGCGGTGGTTGACCAGACAAACC</u>CACATGTCCCACGCTTCCGGGGTCGGCCATCACCACCTCCTTCCGCCACCTAGCGAGCCACCGGTATCTC. The Bxb1 *attB* site is underlined. Transformants were recovered in 2 ml of supplemented 7H9 and shaken overnight at 37 °C. The following day, cells were plated on supplemented 7H10 medium containing 50 μg ml^−1^ hygromycin to select for integration of pKM464 and 2% sucrose for curing of the recombineering plasmid pKM461, which encodes the *sacB* counter-selectable marker. Deletion strains were confirmed by PCR and sequencing of the chromosome-pKM464 junction.

### Analysis of *sigE* mutations in clinical strains

To identify potential PZA resistance signatures of selection on SigE, we contrasted mutation frequency in two sets of isolates. First, a global set of background isolates derived from the Genome-wide Mycobacterium tuberculosis variation database ^68^: PZA-R (n=224) and PZA-S (n =766); and the Global Consortium for Drug-resistant Tuberculosis Diagnostics^69^: PZA-R (n=235) and PZA-S (n=80). Second, a curated set of PZA mono-resistant isolates (n=18). Statistical significance was calculated by Fisher’s Exact Test of independence corrected for multiple hypotheses (Bonferroni method), where the number of non-synonymous mutations observed in *sigE* in any isolate, with a contingency table of categorical variables: “Did/did not have non-synonymous *sigE* mutation” and “Is/is not PZA mono-resistant, PanD_WT_, and PncA_WT_”.

### Evaluation of drug interactions using checkerboard assays

Drug interactions were evaluated through standard checkerboard assays. Briefly, supplemented Middlebrook 7H9 (pH 6.6) was used to culture *M. tuberculosis* H37Rv, *M. tuberculosis* Erdman, or *M. tuberculosis* H37Rv Δ*sigE* to late-log phase. Strains were subcultured to an OD_600_ of 0.01 into 5 ml of supplemented Middlebrook 7H9 (pH 6.6). Bottles were arrayed into rows and columns. PZA was added to each row using a log_2_ dilution scheme from 1600 μg ml^−1^ to 50 μg ml^−1^ and a no drug control. The second drug was subsequently added in a log_2_ dilution scheme to each column. The drug concentration ranges tested against *M. tuberculosis* H37Rv were as follows: meropenem 2 μg ml^−1^ to 0.125 μg ml^−1^, EMB 0.5 μg ml^−1^ to 0.0625 μg ml^−1^, INH 30 ng ml^−1^ to 7.5 ng ml^−1^, D-cycloserine 10 μg ml^−1^ to 1.25 μg ml^−1^. The range for meropenem against *M. tuberculosis* Erdman was 1 μg ml^−1^ to 0.0625 μg ml^−1^. Bottles were incubated at 37 °C and the OD_600_ was measured after 7 days incubation. For checkerboard assays, the MIC was defined as the minimum concentration of drug required to inhibit at least 50% of growth relative to the drug free control. Fractional inhibition concentration index (FICI) was calculated using the following formula: ([MIC of drug B in the presence of a given concentration of Drug A]/[MIC of drug B alone]) + ([MIC of drug A in the of presence of drug B]/[MIC of drug A alone]). If the highest concentration of a drug did not inhibit growth by 50%, the MIC_50_ value used in the FIC calculation was set as 2× the highest concentration that was tested. Drug interactions were defined as follows: FICI values ≤ 0.5 were considered synergistic, FICI values > 0.5 but ≤ 1.0 were considered additive, FICI values from 1.0-4.0 were considered indifferent, and FICI values >4.0 were considered antagonistic^70^.

All relevant data are available upon request.

## Acknowledgements

This study was supported by funds from the National Institutes of Health (AI123146 to A.D.B.). N.A.D was supported by training grant HL007741. R.A was supported by funds from West African Centre for Cell Biology of Infectious Pathogens, University of Ghana (Masters fellowship). S.J.M. and F.V were supported by National Institute of Allergy and Infectious Diseases Grant No. R01AI105185. *M. tuberculosis* strains H37Rv and Erdman, and mycobacteriophage phAE180 were gifts from Dr. William R. Jacobs, Jr. of the Albert Einstein College of Medicine. The H37RvΔ*sigE* and complemented strains were gifts from Dr. Thomas C. Zahrt at the Medical College of Wisconsin. We thank Nicholas D. Peterson for many helpful discussions. We thank Dr. Daryl Gohl of the University of Minnesota Genomics Center for assistance with deep sequencing and data analysis. We thank Jim Werngren and Mikael Mansjö of the Department of Microbiology, Public Health Agency of Sweden, Solna, Sweden for technical help with phenotyping, culturing, and DNA extractions.

## Author contributions

JMT, NAD, MDH, RA, SJM, YM performed experiments; JMT, NAD, MDH, SJM, SEH, FV, YM and ADB analyzed data; JMT, NAD, MDH, SJM, SEH, FV, YM and ADB conceived the work; JMT, NAD, SJM, YM and ADB wrote the manuscript.

